# Integrin and ligand-independent PDGFr signaling synergistically contribute to directional migration of *Xenopus* mesendoderm

**DOI:** 10.1101/318824

**Authors:** Crystal M. Richardson, Bette J. Dzamba, Pooja R. Sonavane, Douglas W. DeSimone

**Affiliations:** Department of Cell Biology, School of Medicine, University of Virginia Health System, P.O. Box 800732, Charlottesville, VA, 22908, USA

## Abstract

Both PDGF signaling and adhesion to fibronectin (FN) matrix have been implicated in the directional collective migration of *Xenopus* mesendoderm cells at gastrulation. However, mesendoderm explants cultured on FN-coated substrates migrate directionally even in the absence of a source of PDGF. Integrin adhesion has been reported to up-regulate PDGF ligand-independent signaling through the PDGF receptor (PDGFr) in cultured mammalian cells. In order to address whether a similar mechanism stimulates PDGFr signaling in the absence of PDGF-A ligand in amphibian mesendoderm, isolated cells were cultured on bacterial fusion proteins containing the Type-III repeats 9-11 of FN (GST-9.11). Type III_9-11_ contains the RGD and “synergy” (PPSRN) sites required for integrin α5β1 adhesion and activation but lacks the PDGF-A ligand-binding site present in the full-length FN protein. In order to ensure mesendoderm was not exposed to PDGF *in vivo* prior to removal and culture *in vitro*, antisense morpholinos were used to inhibit normal expression of PDGF-A ligand in embryos. P-Akt levels were reduced two-fold when either the PDGFr-α was knocked down or when cells were plated on GST-9.11a, which contains a point mutation (PPSRN>PPSAN) that prevents both full activation of integrin α5β1 and cell spreading. Reduced expression of PDGFr-α was accompanied by perturbations in tissue migration, cytoskeletal organization, polarity of cell protrusions, and focal adhesion area. Mesendoderm cells became rounded, and the actin and cytokeratin filaments appeared collapsed and often colocalized near the cell center. Taken together, these findings suggest that integrin adhesion to FN, acting in synergy with PDGFr-α, is sufficient to elevate PI3K-Akt signaling in the mesendoderm even in the absence of the PDGF-A ligand, and to promote forward-directed protrusions and directional tissue migration.

## Introduction

Embryonic development requires precise coordination of cell and tissue movements. How growth factor and adhesion-dependent signaling are integrated during large-scale tissue movements remains unclear. Morphogens, chemokines, and soluble growth factors guide directional movement (Haeger et al., 2015). Platelet derived growth factor (PDGF) is important for single cell chemotaxis and directional collective cell migration in many systems. Cells undergo chemotaxis toward PDGF during wound healing (Lynch et al., 1987; Schneider et al., 2010), cancer invasion (Andrae et al., 2008; Watts et al., 2016; Yeh et al., 2016), *Drosophila* border cell migration (Duchek et al., 2001; McDonald et al., 2003), *Xenopus* mesoderm migration (Nagel et al., 2004; Smith et al., 2009; Symes et al., 2010), and zebrafish mesendoderm migration (Montero et al., 2003). PDGF receptor (PDGFr) expression correlates with advanced tumor stages and metastasis in some cancers (Andrae et al., 2008). Integrin-FN adhesive signaling can activate PDGFr independent of the PDGF ligand (Sundberg and Rubin, 1996; Veevers-Lowe et al., 2011). Although it is clear that PDGF signaling is important for directional migration (Duchek et al., 2001; Nagel et al., 2004; Yang et al., 2008), whether ligand independent signaling through the PDGFr is sufficient to promote such movement remains unclear.

The migration of *Xenopus* mesendoderm cells at gastrulation is an example of a collective cell movement driving tissue morphogenesis (Winklbauer, 1990). PDGF-A ligand in this system is reported to act as a directional cue to orient cell protrusions in the direction of the animal pole of the blastocoel roof (BCR) (Nagel et al., 2004; Smith et al., 2009). PDGF-A ligand is expressed by the BCR cells in two alternatively spliced forms: a short form and a long form, which are both secreted at gastrulation. The short form is freely diffusible and acts a long-range signal for radial intercalation of prechordal mesoderm cells toward the ectoderm (Damm and Winklbauer, 2011). The long form has an ECM-binding domain that allows PDGF-A ligand to attach to the HepII region of FN (Smith et al., 2009). Mesendoderm cells express the corresponding PDGF receptor α (PDGFr-α). As these cells migrate on the BCR, they contact assembled FN matrix with sequestered PDGF-A ligand (Ataliotis et al., 1995; Damm and Winklbauer, 2011). Leader cells of the mesendoderm are the first to contact this matrix via integrin based adhesion and send out forward directed protrusions (Nagel et al., 2004). The importance of PDGF as a major guidance cue required for directional mesendoderm migration has been explored using BCR-conditioned substrates (Nagel et al., 2004; Smith et al., 2009; Symes et al., 2010). Knockdown of the PDGF-A ligand results in misdirected protrusions on BCR conditioned substrates, suggesting a role for PDGF-A ligand bound to FN in chemotaxis (Nagel et al., 2004).

However, directional migration of explanted mesendoderm tissue is reported to occur on nonfibrillar FN substrates in the absence of matrix-attached PDGF-A ligand (Davidson et al., 2002; Weber et al., 2012), calling into question the importance of PDGF-A ligand in this process. One possible explanation is that ligand-independent activation of the PDGFr is responsible for directed migration of explanted mesendoderm. Thus, we sought to investigate the relationship between PDGFr dependent signaling and integrin engagement of FN in directional migration. This study reports that PDGFr signaling can activate PI3K-Akt to regulate the organization of cytoskeleton to orient protrusions in the absence of PDGF-A ligand.

While growth factor receptors typically become activated as a result of direct molecular interactions with their cognate ligands, growth factor receptors can also be activated in a ligand independent manner. For example, the PDGFr is phosphorylated at Tyr-751 and activated following cell adhesion to FN resulting in integrin α5β1 activation and enhanced downstream signaling leading to the phosphorylation of Fak at Tyr-397 (Veevers-Lowe et al., 2011). This crosstalk between integrin α5β1 and the PDGFr may initiate a positive signaling feedback loop, in which FAK phosphorylates and activates the PDGFr (Veevers-Lowe et al., 2011). Ligand independent activation of EGFr occurs in lung cancer, glioblastoma, and squamous cell carcinoma (Guo et al., 2015; Shen and Kramer, 2004). Integrin β1 and E-cadherin associate with and cause the activation of EGFr in a ligand independent manner. (Moro et al., 1998; Shen and Kramer, 2004). In each of these examples, growth factor signaling is up-regulated through cooperation with cell adhesion molecules. However, the interplay between growth factor receptor and integrin signaling during morphogenesis remains unclear. The current study explores the relationship between PDGF and integrin signals to promote cell adhesion and directional lamellipodial protrusion formation in the collective cell migration of *Xenopus* mesendoderm.

## Results

### PDGFr-α and integrin-fibronectin (FN) adhesions contribute to Akt phosphorylation

Mesendoderm cells express integrin α5β1, which binds to FN RGD in cooperation with the synergy site sequence (Ramos and DeSimone, 1996) resulting in integrin activation and transition to a “high affinity” state (Li et al., 2003). Integrin α5β1 binding to RGD containing Type III_10_ repeat alone is sufficient for integrin adhesion to FN although the integrins remain in a “low-affinity” state (García et al., 2002). Cells can attach to RGD-containing fragments of FN but are unable to spread when the synergy site in Type III_9_ is mutated (Ramos and DeSimone, 1996). Integrin binding to FN can lead to the activation of the PDGFr in a PDGF ligand independent manner (Sundberg and Rubin, 1996; Veevers-Lowe et al., 2011) and this upregulation of signaling can further stimulate integrin conformational change and activationVeevers-Lowe et al., 2011). Inputs from both integrin-FN adhesions and PDGF signaling mediate cell attachment and cell spreading (Ramos and DeSimone, 1996; Symes and Mercola, 1996). Although it is clear that PDGF signaling and integrin adhesive signaling act cooperatively, the relative importance of PDGFr-α dependent signaling and integrin signaling remains unclear.

To investigate whether PDGFr-α regulates integrin dependent adhesive signaling in directional *Xenopus* mesendoderm migration, an antisense morpholino oligodeoxynucleotide (PDGFr-α MO) was designed to block the translation of *Xenopus* PDGFr-α mRNA. PDGFr-α MO mesendoderm cells were plated on purified FN fusion proteins (Fig. 1 A). Bacterially purified FN fusion proteins were used instead of purified plasma FN to eliminate the possibility of pFN contamination with PDGF ligand. Moreover, using FN fusion proteins allows for the evaluation of the contribution of the FN RGD and synergy sites to cell adhesion and cell signaling (Li et al., 2003; Ramos et al., 1996). The FN fragments used in these experiments correspond to Type-III repeats 9 through 11 of *Xenopus* FN containing the RGD sequence and the synergy site (PPSRN), which are the key adhesion sites for full integrin α5β1 engagement on FN (Fig. 1 A). To assess the contribution of integrin engagement with FN synergy site to signaling, the 9.11 fragment which has a point mutation in the synergy site, 9.11a, was used (Fig. 1 A). As previously demonstrated Control morphant (Control MO) mesendoderm cells on 9.11 attach and spread whereas cells on 9.11a attach but are unable to spread (Fig. 1 B) (Ramos et al., 1996). Decreased cell spreading was also noted in PDGFr-α MO-treated cells on 9.11 compared to Control MO cells (Fig. 1 B, 24% decrease). These data are consistent with a role for both integrin engagement with FN and PDGFr-α functioning in mesendoderm cell spreading on FN.

**Figure 1.**
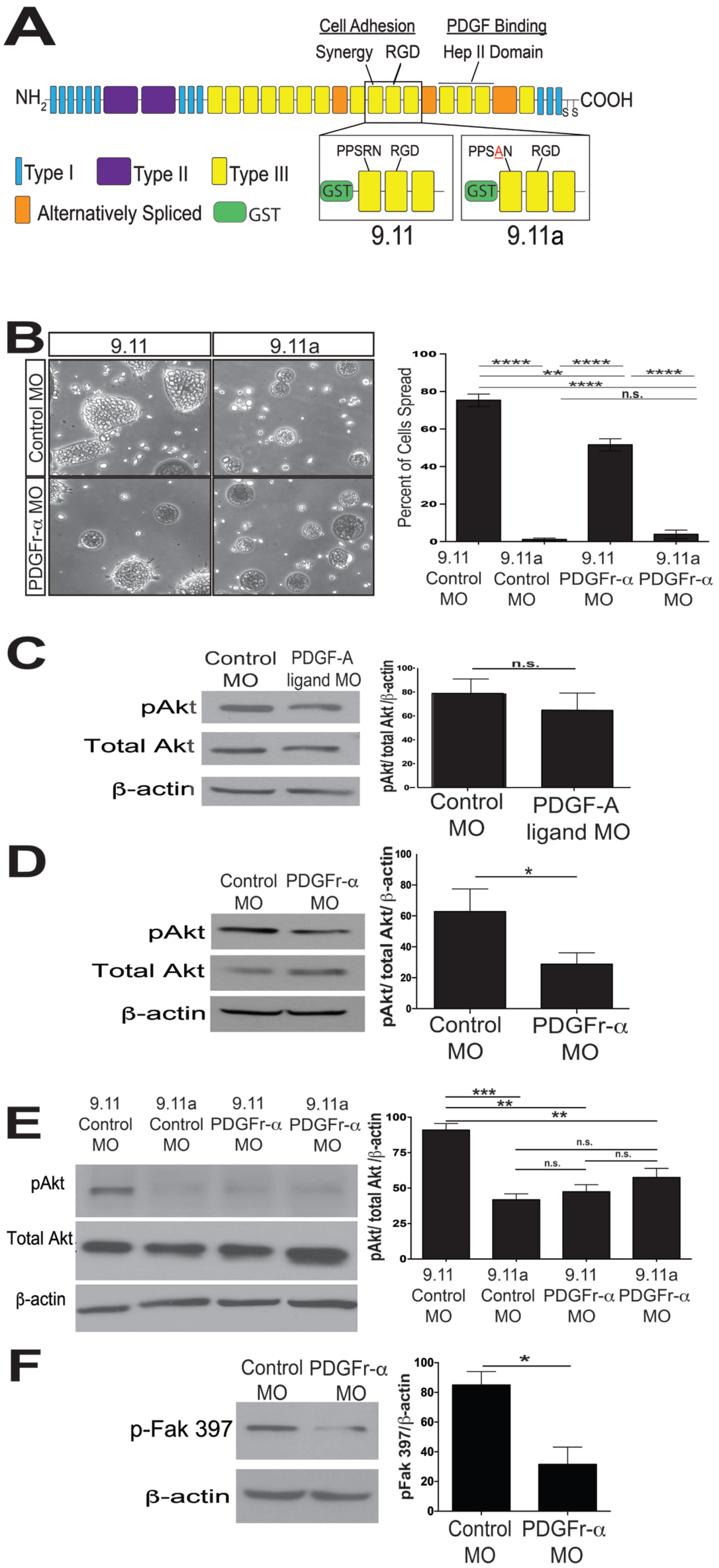
PDGFr-α and integrin adhesion contribute to Akt phosphorylation. (A) Diagram of full-length *Xenopus* fibronectin (FN) and FN bacterial GST fusion proteins 9.11 and 9.11a. 9.11 and 9.11a contain FN Type III repeats 9-11 including the RGD site in Type III_10_ “synergy” site (PPSRN) in Type III_9_; 9.11a has a point mutation in the synergy site (PPS**A**N)that inhibits integrin activation. Type I repeats are light blue, Type II repeats are purple, and Type III repeats are yellow. (B) Representative images of Control MO and PDGFr-α MO mesendoderm cells on 9.11 and 9.11a with corresponding quantification of cell spreading. Number of cells analyzed: 9.11 Control MO = 571 cells, 9.11A Control MO = 393 cells, 9.11 PDGFr-α MO = 586 cells, and 9.11A PDGFr-α MO = 402 cells across 3 separate experiments. (C) Western blot of p-Akt levels in control and PDGF-A ligand MO Stage 11 embryos. Graph is (*N* = 3). (D) Western blot of p-Akt levels in control and PDGFr-α MO gastrula Stage 11 embryos. Graph is (*N* = 8). (E) Western blot of p-Akt levels of Control MO and PDGFr-α MO mesendoderm cells plated on 9.11 or 911a. Graph is (*N* = 3). (F) Representative Western blot of p-Fak 397 for Control MO and PDGFr-α MO gastrula Stage 11 embryos. Graph is (*N* = 3). (B-F) Data are mean + standard error of the mean. A single asterisk (*) indicates *p* <.05, a double asterisk (**) indicates *p* <.005, a triple asterisk (***) indicates *p* <.001, and n.s. indicates no statistically significant difference.

It is well established that both α5β1 integrin-FN dependent adhesive signaling (King et al., 1997) and PDGF activate PI3K-Akt signals (Franke et al., 1995). However, it has only recently been demonstrated that α5β1 integrin-FN dependent adhesive signaling and PDGFr can act synergistically to enhance PI3K-Akt signals (Veevers-Lowe et al., 2011). To evaluate the relative contribution of integrins and PDGFr-α to the phosphorylation of Akt (p-Akt), Western blot analysis was performed comparing Control MO, PDGF-A ligand MO, and PDGFr-α MO-treated embryos. When the PDGF-A ligand is knocked down, the levels of p-Akt were not significantly different from Control MO (Fig. 1 C), whereas PDGFrα knockdown results in a significant reduction of pAkt levels (Fig. 1 D). Because there was a decrease in p-Akt levels when the PDGFr-α was knocked down, but not when the PDGF-A ligand was knocked down, the role of ligand independent PDGFr-α signaling in the phosphorylation of Akt was further evaluated using FN fusion proteins lacking the PDGF-A ligand binding site.

A 50% decrease in p-Akt levels was noted in Control MO-treated cells on 9.11a compared to cells on 9.11 (Fig 1 E). Knocking down the PDGFr-α also caused a 50% decrease in p-Akt levels, similar to decreased p-Akt levels of Control MO cells on 9.11a (Fig. 1 E). When PDGFr-α was knocked down in conjunction with disrupting integrin adhesive signals by plating cells on 9.11a, there was no additive effect and no additional decrease in p-Akt was noted (Fig. 1 E). Thus reduced cell spreading correlated with decreased p-Akt levels. In order to maintain full p-Akt levels and cell spreading, both the FN synergy site and the PDGFr-α are required. These data are consistent with crosstalk between PDGFr-α and integrin-FN adhesive signals leading to cell spreading and the phosphorylation of Akt.

Because PDGF and integrin dependent signaling converge on the downstream targets FAK, PI3K, and AKT (Sieg et al., 2000; Reif et al., 2003; Higuchi et al., 2012) whether PDGFr-α dependent signals were important for the phosphorylation of FAK at Tyr 397 was next investigated (Fig. 1 F). An approximately 50% decrease in levels of p-Fak at Tyr 397 in PDGFr-α MO embryos was noted (Fig. 1F). This decrease correlated with the decrease in p-Akt levels (Fig. 1 D). Taken together, these data are consistent with PDGFr-α functioning in cooperation with integrin adhesive signaling to activate downstream signaling pathways that lead to the phosphorylation of Akt and Fak at Tyr 397 to regulate cell spreading on FN.

### PDGFr-α stabilizes mesendoderm cell adhesions with FN

Because mesendoderm explants migrate directionally along nonfibrillar FN without added PDGF-A (Davidson et al., 2002), the functional importance of PDGFr dependent signaling in directional mesendoderm migration is unclear. To test the role of PDGFr signals in mesendoderm migration, the leading edge of the mesendoderm was tracked over time. Migration of Control MO, PDGF-A ligand MO, and PDGFr-α MO treated mesendoderm explants were compared. Mesendoderm is not known to express PDGF-A ligand (Ataliotis et al., 1995; Damm and Winklbauer, 2011) and PDGF was not added to explant culture, but to reduce the possibility that the mesendoderm was exposed to PDGF-A *in vivo* prior to explantation, explants were prepared from PDGF-A MO injected embryos. The PDGF-A MO construct was used previously and caused disruption of directional mesoderm migration on blastocoel roof conditioned substrates that contain FN bound PDGF-A ligand (Nagel et al., 2004). The contribution of PDGFr-α to mesendoderm migration on FN in the absence of PDGF has not yet been evaluated.

When the leading edge of the mesendoderm tissue was tracked over 30 minutes a significant increase in retractions in PDGFr-α MO explants was noted compared to Control MO explants (Fig. 2 A). Retractions were defined as any rearward movement opposite the direction of migration. A significant decrease in the distance traveled was observed in PDGFr-α MO mesendoderm explants compared to Control MO explants at all timepoints after 20 minutes (Fig. 2 B). However, there was no significant difference in distance traveled between PDGF-A ligand MO explants and Control MO explants noted at any timepoint (Fig. 2 B).

**Figure 2.**
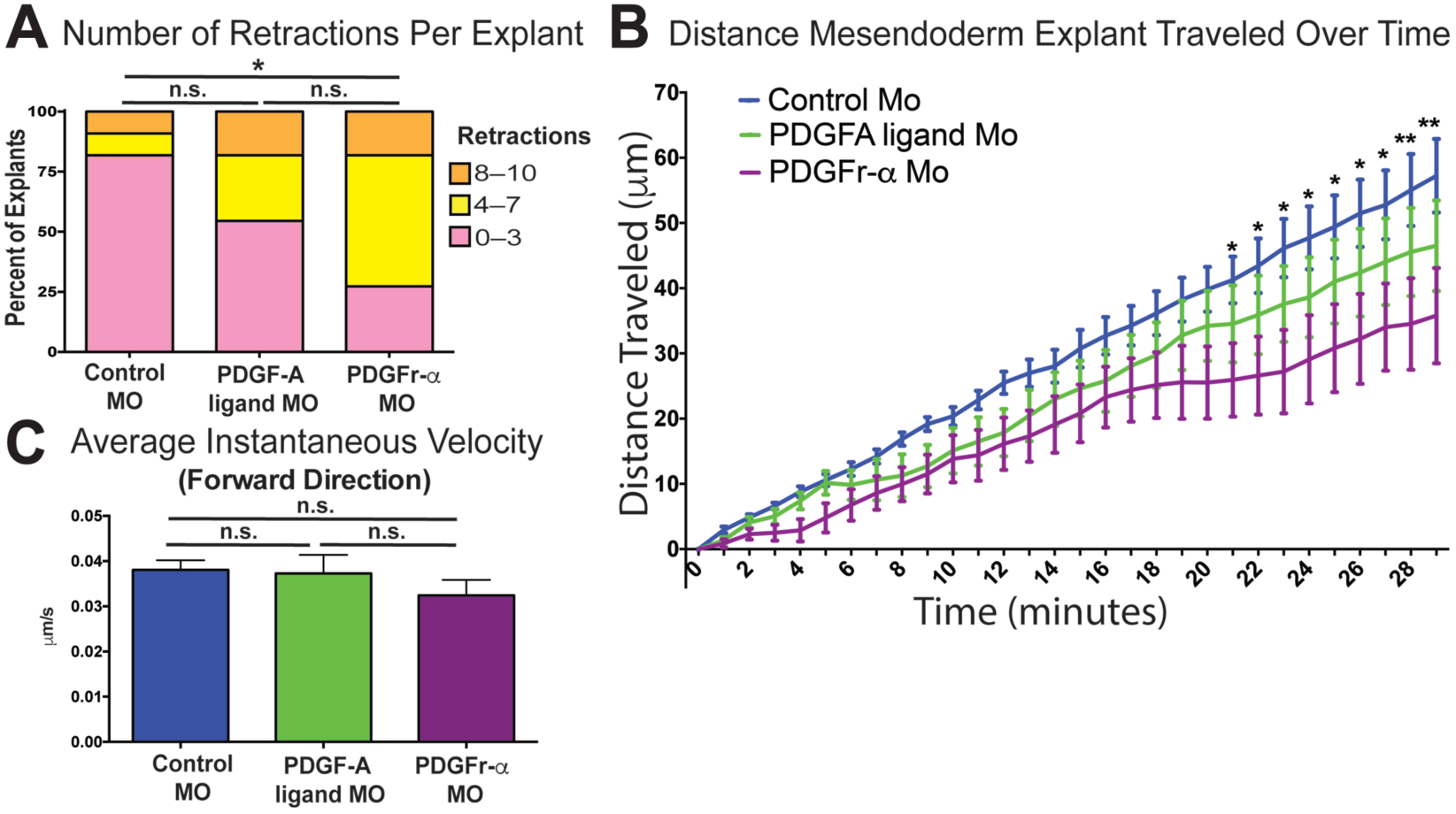
Loss of PDGFr-α increases retractions in mesendoderm explants. (A) Quantification of the number of times mesendoderm explants retracted over a 30-minute time period. Data represent the percentage of explants with high (8–10, orange bar), medium (4–7, yellow bar), and low (0–3, pink bar) numbers of retractions. (B) Quantification of the average distance mesendoderm explants traveled over a 30-minute time period. (A–B) A single asterisk (*) indicates *p* <.05 and a double asterisk (**) indicates *p* <.005. (B) No statistically significant differences were noted between Control MO and PDGF-A ligand MO explants at any timepoint. (C) Quantification of the average instantaneous velocity calculated in the forward direction, excluding retractions. n.s. indicates no statistically significant difference. (B-C) Data are represented as mean + standard error of the mean. (A–C) 11 individual explants analyzed per treatment across 3 separate experiments.

In order to determine if the observed decrease in the distance traveled by PDGFr-α morphant mesendoderm tissue was primarily due to the increased retraction of the tissue we next investigated whether PDGF signaling influences the migration rate of mesendoderm. The instantaneous velocity was measured in Control MO, PDGF-A ligand MO, and PDGFr-α MO mesendoderm explants. There was no significant change in the instantaneous velocity of mesendoderm explants when either the PDGFr-α or the PDGF-A ligand was knocked down compared to controls (Fig. 2 C). These data are consistent with increased retractions resulting in a decrease in the distance of the mesendoderm tissue migration over time but not a reduction in migration speed *per se*.

### PDGFr-α knockdown disrupts monopolar protrusive cell behavior

Because integrin-dependent adhesive signaling is required for cell protrusion formation and PDGF has been implicated in orienting cell protrusions during migration, we next evaluated the contribution of PDGF signaling to regulating polarized protrusive cell behavior during mesendoderm migration. To test whether PDGFr-α was necessary for forward directed lamellipodial formation, protrusion angles were quantified relative to a line parallel to the direction of explant travel through the center of the cell body. Measured protrusion angles were binned in 30° increments and the percent of protrusions in each bin was plotted in rose diagrams. 180° represents the protrusions in the direction of travel (Fig. 3). The protrusion angles provide a measure of protrusive behaviors that are characteristic of migrating mesendoderm (Bjerke et al., 2014; Davidson et al., 2002). A 34% decrease in forward directed protrusions along the leading edge was noted in PDGFr-α MO explants compared to Control MO explants (Fig. 3 A -F). The percent of forward directed protrusions in PDGF-A ligand MO explants was not significantly different from Control MO explants (Fig. 3 D -E). These results indicate a role for the PDGFr-α in maintaining forward directed protrusions during mesendoderm migration.

**Figure 3.**
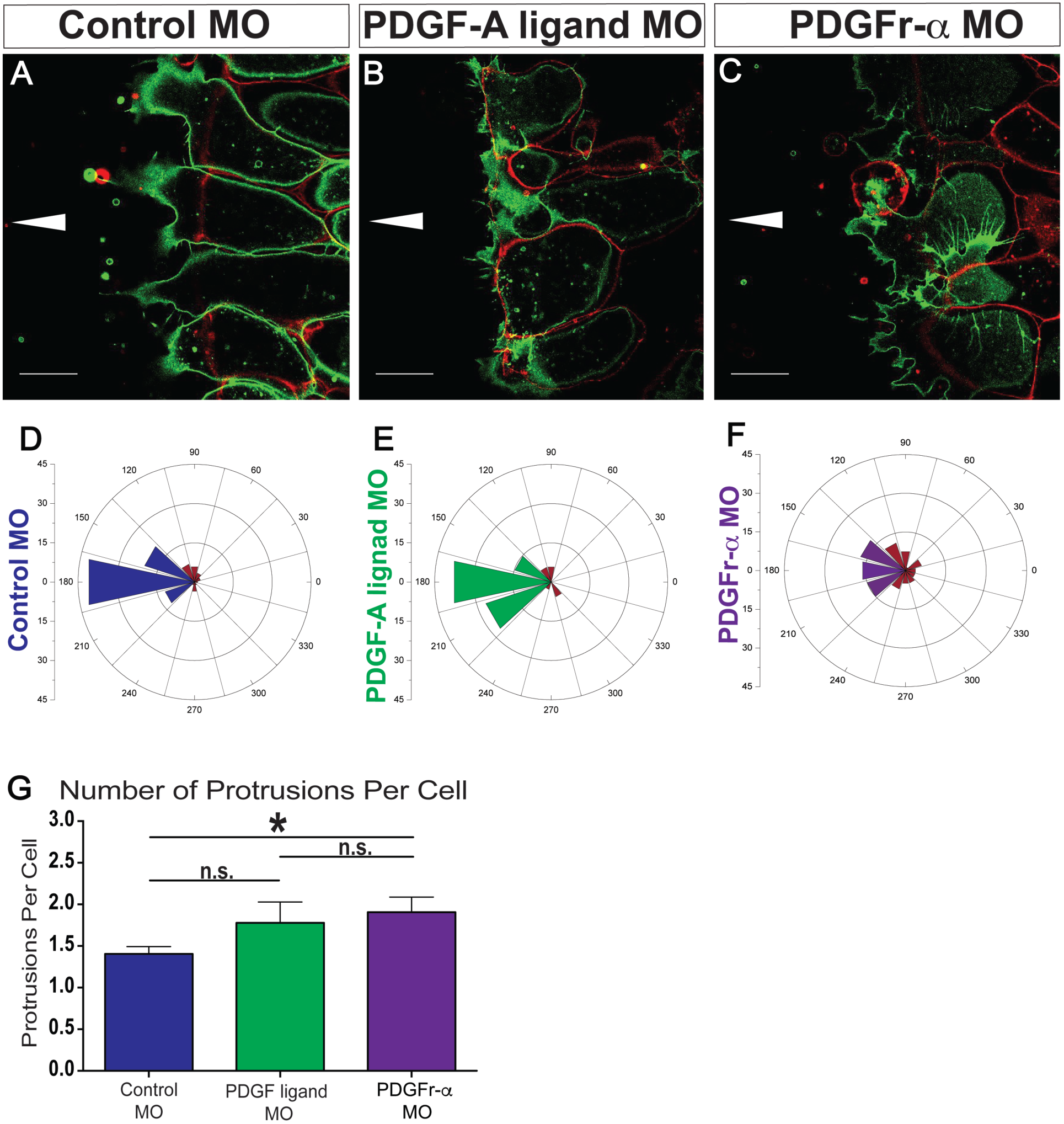
PDGFr-α is required for directional monopolar protrusive activity. (A–C) Representative confocal images of live Control MO, PDGF-A ligand MO, and PDGFr-α MO mesendoderm explants migrating on FN and expressing membrane-EGFP. Overlay of images taken at the plane of the coverslip (pseudocolored green) and 5μm above (pseudocolored red) to simultaneously visualize the cell body at 5-μm above substrate and cell protrusions at the substrate. (D–F) Rose diagrams representing cell protrusion angles. Protrusion angles are measured relative to the cell centroid and plotted with the center of the rose diagram representing the cell centroid. An angle of 180° represents the direction of migration, and angles within 150°– 210° are defined as normally oriented protrusions. Angles outside 150°–210° are defined as misdirected and are pseudocolored red in the rose diagram. (G) Quantification of the average number of protrusions per cell. Data are mean + standard error of the mean. A single asterisk (*) indicates *p* <.05 and n.s. indicates no statistically significant difference. (D) Number of protrusions analyzed: Control MO = 84, PDGF-A ligand MO = 34, and PDGFr-α MO = 61 from 7 experiments.

Migrating mesendoderm cells display monopolar protrusive behavior, cells typically extend 1-2 protrusions in the direction of travel (Bjerke et al., 2014; Davidson et al., 2002). To determine whether PDGFr-α was important for this monopolar protrusive behavior, the number of protrusions per cell was counted in migrating mesendoderm explants. An increase in the average number of protrusions per cell was noted when PDGFr-α was knocked down, whereas the average protrusion number in PDGF-A ligand MO cells was not significantly different from Control MO cells (Fig. 3 G).

### PDGFr-α functions in lamellipodial protrusion formation of the actin cytoskeleton

PDGF signaling has been shown to lead to the loss of stress fibers in 3T3 cells (Herman and Pledger, 1985; Nagano et al., 2006). In order to assess whether PDGFr-dependent signaling alters actin organization in mesendoderm we used phalloidin to visualize the actin cytoskeleton. Thick actin-filled lamellipodial protrusions oriented in the direction of migration were observed in Control MO and PDGF-A ligand MO explants (Fig. 4 A, red arrowheads). However, the lamellae were reduced in PDGFr-α MO mesendoderm compared to controls (Fig. 4 A-C). Protrusions were more filopodia-like in appearance with reduced size and misdirected in PDGFr-α MO mesendoderm explants (Fig. 4 C). A fine actin filament network extended throughout the cell in Control MO explants (Fig. 4 A). Actin structures appeared collapsed and cells had a decrease in fine actin filaments that extended throughout the cell in PDGFr-α MO explants (Fig. 4 C). Traces of actin-based protrusions were observed in front of the mesendoderm tissue when PDGFr-α was knocked down (Fig. 4 C, blue arrowheads). A possible explanation for this is that the cell membrane and actin were left on the FN glass coverslip after tissue retraction. These data are consistent with decreased integrin adhesive signaling and reduced p-Akt (Fig. 1) leading to mesendoderm tissue retraction and cells rounding up in PDGFr-α MO mesendoderm explants (Fig 3 C).

**Figure 4.**
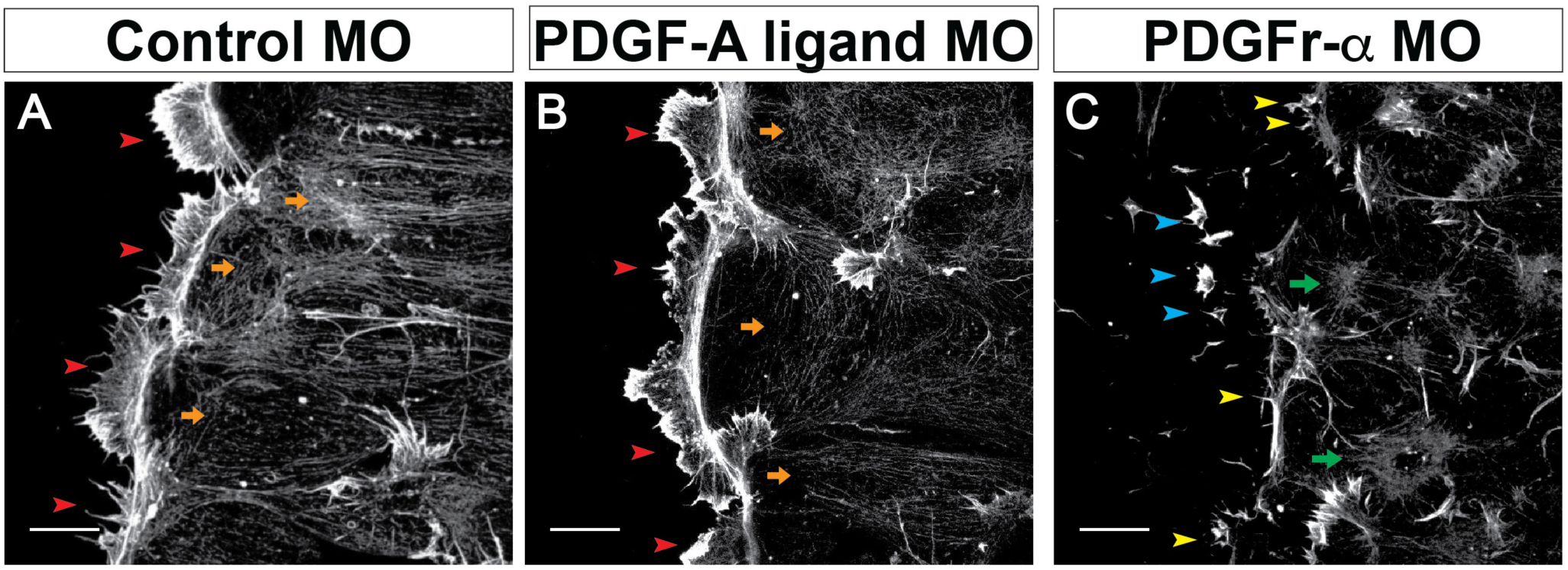
Actin organization is altered in PDGFr-α MO explants. (A–C) Representative collapsed 20 μm z-stack confocal images of fixed mesendoderm explants stained with phalloidin to visualize F-actin. (A–B) Broad actin-rich lamellipodial protrusions at the leading edge of Control MO and PDGF-A ligand MO explants (red arrowheads). Fine F-actin filaments that extend to the edges of the mesendoderm cells (orange arrows). (C) Fragments of membrane containing F-actin found in the front of the PDGFr-α MO mesendoderm explant (blue arrowheads). “Filopodia-like” protrusions (yellow arrowheads) and collapsed F-actin in PDGFr-α MO explants (green arrows). (A–C) Representative images from a total of : Control MO = 11, PDGF-A ligand MO = 13, and PDGFr-α MO = 11 from 4 experiments. Scale bars are each 20μm.

### Cytokeratin intermediate filament network collapses in PDGFr-α MO mesendoderm explants

Mesendoderm monopolar protrusive behavior is dependent on the organization of the keratin intermediate filament cytoskeleton (Weber et al., 2012) Because the orientation and number of mesendoderm cell protrusions was disrupted in PDGFr-α MO explants (Fig. 3), cytokeratin cytoskeletal organization was next investigated. Cytokeratin networks extended throughout the cells in Control MO (Fig. 5 A, D, G) and PDGF-A ligand MO (Fig. 5 B, E, H) explants, however cytokeratin appeared to collapse to the center of the mesendoderm cells in PDGFr-α MO explants (Fig. 5 C, F, I). Collapses in the cytokeratin network overlapped with actin foci in PDGFr-α MO explants (blue arrows in Fig. 5 F, I, L).

**Figure 5.**
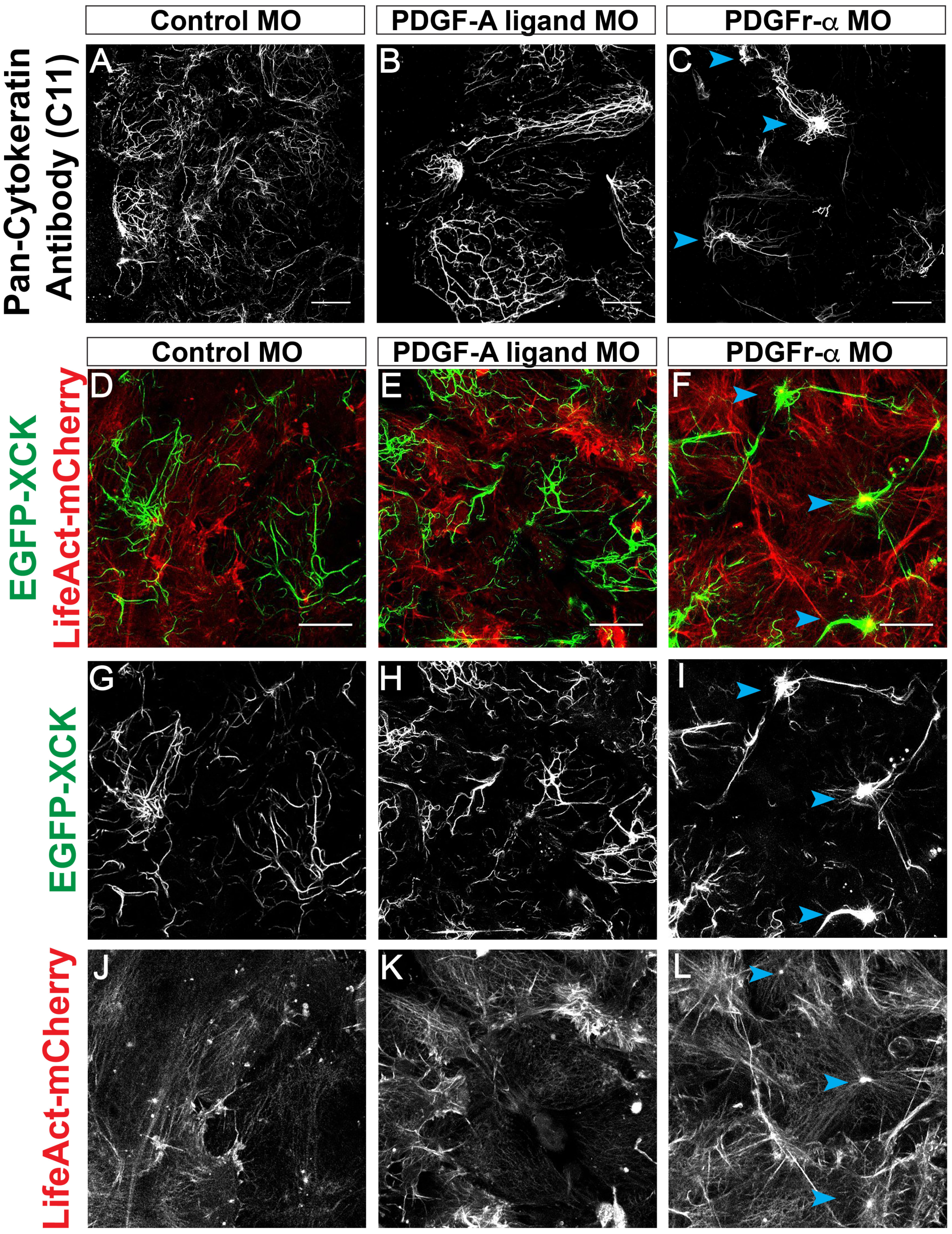
Changes in actin organization are accompanied by disruptions in the cytokeratin intermediate filament network in PDGFr-α MO explants. (A–C) Representative collapsed z-stack confocal images of fixed mesendoderm explants stained with pan-cytokeratin antibody (C11) to visualize the cytokeratin intermediate filament network. Number of explants: Control MO = 24, PDGF-A ligand MO = 12, and PDGFr-α MO = 10 from 7 experiments. (D–L) Representative collapsed z-stack confocal images of mesendoderm explants expressing EGFP-*Xenopus* cytokeratin (EGFP-XCK) and LifeAct-mCherry. EGFP-XCK is pseudocolored green, and LifeAct-mCherry is pseudocolored red. Number of explants: Control MO = 8, PDGF-A ligand MO = 4, and PDGFr-α MO = 9 from 3 experiments. Blue arrowheads indicate cytokeratin filaments that are collapsed in (C, F, I, L). Single-channel images of EGFP-XCK (G-I) and LifeAct-mCherry (J–L). Scale bars are 20 μm.

### PDGFr-α MO mesendoderm cells have larger focal adhesions

The increase in tissue retractions (Fig. 2) and decrease in p-Fak at Tyr 397 in PDGFr-α MO explants (Fig. 1F), led us to ask whether focal adhesion size was altered in PDGFr-α MO explants. To visualize focal adhesions, paxillin-GFP was expressed and imaged in migrating mesendoderm explants using TIRF microscopy (Fig. 6 A-L). A significant increase in the size of focal adhesions was noted upon disruption of PDGF signaling with the largest focal adhesions in PDGFrα MO-treated explants (Fig. 6 L-M). Our data fits with the current understanding of Fak regulation of focal adhesion size. Increased focal adhesion size has also been reported in mouse embryonic mesodermal Fak (-/-) cells (llić et al., 1995).

**Figure 6.**
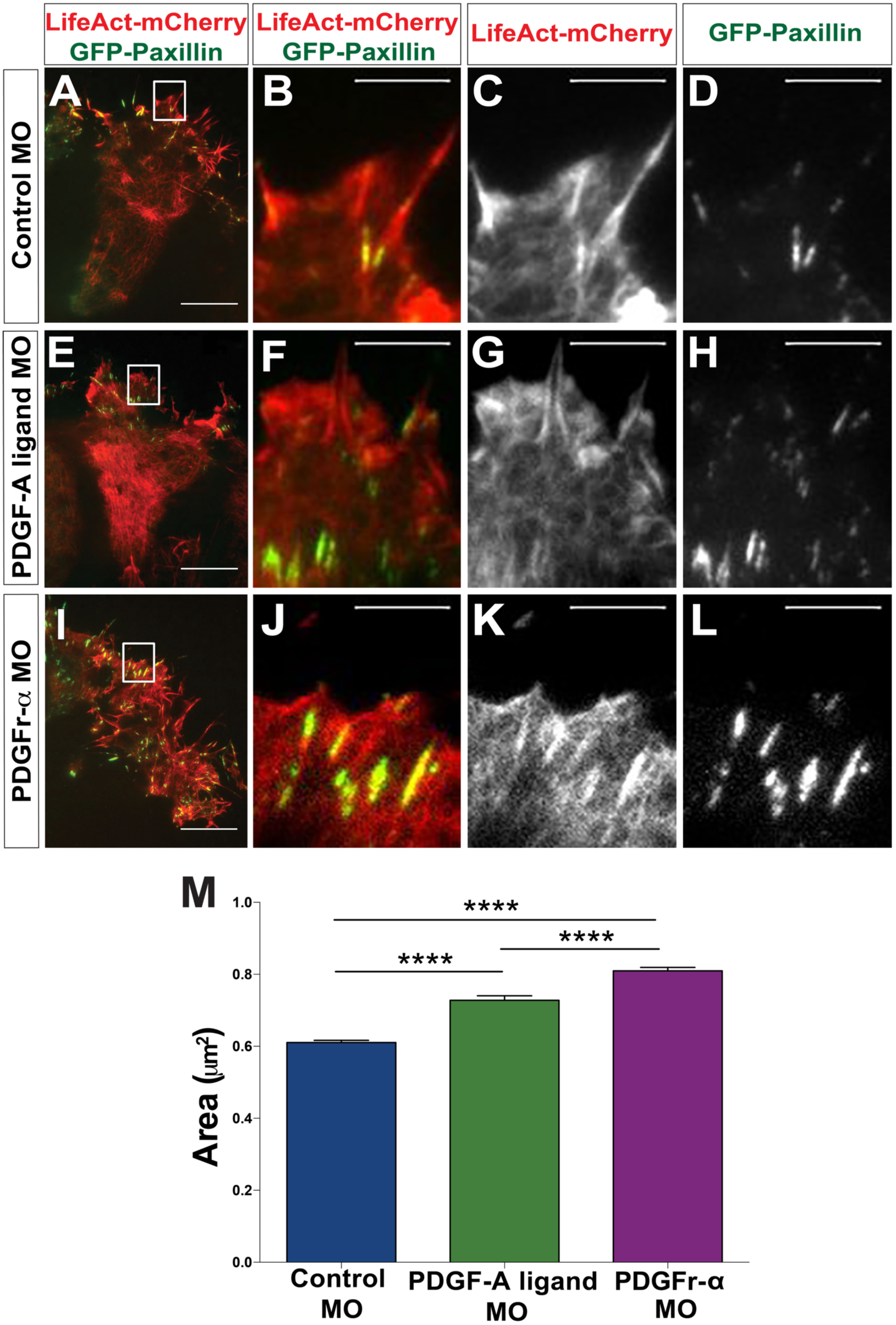
Focal adhesion area is increased in PDGFr-α MO explants. Mesendoderm explants expressing LifeAct-mCherry, pseudocolored red, and Paxillin-GFP, pseudocolored green. Live mesendoderm explants were imaged using TIRF microscopy to visualize paxillin-rich focal adhesions on the fibronectin substrate. (A, E, I) White boxes indicate regions enlarged in panels (B–D, F–H, J–L). (A–D) Control MO explants. (E–H) PDGF-A ligand MO explants. (I–L) PDGFr-α MO explants. (M) Quantification of focal adhesion area. Data are mean + standard error of the mean. *p* <.001 for each comparison made. Control MO (*N* = 5, n=14 explants), PDGF-A ligand MO (*N* = 3, n=7 explants), PDGFr-α MO (*N* = 5, n=16 explants)

## Discussion

### Integrin and PDGF signals are required for directional *Xenopus* mesendoderm migration

Integrins are important mediators of cell signaling and are required for cell adhesion to the surrounding extracellular matrix. Integrin receptors can function in conjunction with other receptors, such as growth factor receptors, to enhance downstream cell signaling pathways (Ross, 2004; Veevers-Lowe et al., 2011; Ivaska and Heino, 2012;Brizzi et al., 2012). This study provides insight into the interplay between integrin and growth factor signaling during embryonic development. Here, a role for integrin signaling in cooperation with PDGF signaling was identified to regulate cytoskeletal dynamics, focal adhesion assembly, and to orient cell protrusions during directional migration of *Xenopus* mesendoderm. This model builds on previous studies that suggested PDGF-A ligand acts as a chemoattractant to enhance directional migration (Ataliotis et al., 1995; Nagel et al., 2004; Smith et al., 2009; Symes and Mercola, 1996). We conclude that PDGFr-α can synergize with integrin-fibronectin adhesive signaling to direct mesendoderm migration.

Although it is known that integrins and growth factors converge to activate the PI3K-Akt signaling pathway (Baron et al., 2002; Baron et al., 2003; Veevers-Lowe et al., 2011; Higuchi et al., 2012, how integrins synergize with growth factors remains unclear. The current study identified functions for the PDGFr-α in the absence of PDGF-A ligand. The PDGFr-α MO phenotype has features in common with disruption of integrin-FN adhesions using function blocking antibodies (Davidson et al., 2002; Ramos and DeSimone, 1996; Winklbauer and Nagel, 1991) or knocking down Fak (Bjerke et al., 2014). Inhibiting the interactions of α5β1integrin with the RGD site results in mesendoderm tissue explants detaching and “snapping-back” (Davidson et al., 2002). Fak MO cells are less spread on FN but explants do not detach (Bjerke et al., 2014). This phenotype is comparable to cells on 9.11a that are unable to attach to the FN “synergy site” (Bjerke et al., 2014; Davidson et al., 2002; Ramos et al., 1996). Similar to Fak MO and cells on 9.11a, PDGFr-α MO cells are less well spread on FN. Although PDGFr-α MO mesendoderm explants are able to migrate on nonfibrillar FN substrates, there is an increase in the number of retractions compared to Control MO explants and this is consistent with a defect in integrin-fibronectin adhesions or a defect in the cytoskeleton. Our study also demonstrated a role for integrin adhesive signaling in cooperation with PDGFr-α dependent signaling in the phosphorylation of Akt at Tyr 308. Taken together, these findings increase our understanding of how integrin and PDGF signals enhance PI3K-Akt signals to produce collective cell migration of *Xenopus* mesendoderm tissue.

### Ligand independent signaling of the PDGFr

The PDGF ligand is a well-established chemoattractant that promotes the orientation of cell protrusions to direct cell migration (Lynch et al., 1987; McDonald et al., 2003; Montero et al., 2003). *Xenopus* mesoderm can respond to PDGF-A ligand embedded in the FN of blastocoel conditioned substrates by orienting protrusions in the direction of the animal pole (Nagel et al., 2004). An alternatively spliced form of the PDGF-A ligand that lacks the C-terminal matrix-binding domain is freely diffusible and unable to bind FN or promote directional mesendoderm migration (Nagel et al., 2004). Although it remains unclear whether a protein gradient of the PDGF-A ligand forms in the developing embryo, it is clear that the FN-bound form of PDGF-A ligand can function as a chemoattractant to enhance directional mesendoderm migration (Nagel et al., 2004).

Directional mesendoderm migration occurs *in vitro* when explants that contain the entire dorsal marginal zone of early gastrula stage embryos including ectoderm are plated on unmodified nonfibrillar FN without attached PDGF-A ligand (Davidson et al., 2002). The mesendoderm in these multi-tissue explants becomes polarized in response to mechanical cues (Weber et al., 2012). This calls into question the role of PDGF as a FN-attached chemotactic cue in mesendoderm migration. If mesendoderm migration is directional in the absence of PDGF-A ligand, then is PDGF signaling required for directional migration? Integrin-FN adhesive signaling can activate the PDGFr (Sundberg and Rubin, 1996; Veevers-Lowe et al., 2011). Does ligand independent PDGFr signaling enhance directional *Xenopus* mesendoderm migration? And if so, how do PDGFr dependent signals act in synergy with other cues to direct migration?

The current study sought to address these questions using a morpholino approach to knock down PDGFr-α. This study demonstrated that PDGFr-α functions to promote directional migration during mesendoderm migration on FN. Interestingly, PDGFr-α functions in the absence of a gradient of sequestered PDGF-A ligand. These data support a role for integrin-fibronectin adhesive signals acting in cooperation with the PDGFr-α. In the developing embryo, integrin cooperation with PDGFr-α may provide another way to enhance directional migration. One possibility is maintaining a ligand gradient over long-range migration can be difficult to control spatiotemporally. Highest traction stresses and focal adhesions are localized to the front of the migrating mesendoderm explants (Sonavane et al. 2017), supporting a model where integrins are in the highest activation state in the front of the tissue where they spatially enhance PDGFr dependent signaling.

### PDGFr functions to maintain the organization of the underlying cytoskeleton and normal focal adhesion size

Actin-filled lamellipodial protrusions are a prominent feature of directional *Xenopus* mesendoderm migration (Bjerke et al., 2014; Davidson et al., 2002). PDGF signaling alters the actin cytoskeleton leading to a reduction of stress fibers in 3T3 cells (Herman and Pledger, 1985; Nagano et al., 2006). We report that ligand independent signaling by the PDGFr-α is necessary to maintain actin-filled lamellipodial protrusions. In the absence of the PDGFr-α protrusions become more “filopodial-like” and become mis-directed. The normal arrangement of the cortical actin cytoskeleton including fine actin filaments found throughout the cell body is dependent on PDGFr-α. The collapse in the actin cytoskeletal network observed in PDGFr-α MO mesendoderm explants likely contributes to the aberrant migratory behavior.

The organization of both the actin and cytokeratin intermediate filament cytoskeleton is dependent on the PDGFr-α. Recruitment of keratin intermediate filaments to C-cadherin junctions is a result of asymmetric tissue tension and keratin is required for directional protrusion formation (Weber et al., 2012). Knockdown of either plakoglobin or Fak results in a disruption in the cytokeratin intermediate filament network without significantly altering cell cohesion within the mesendoderm tissue (Bjerke et al., 2014; Weber et al., 2012). Similarly, knockdown of the PDGFr-α failed to disrupt cell-cell contacts but recruitment of keratin intermediate filaments to cell contacts was disrupted. Keratin intermediate filaments become collapsed toward the center of the cell where they co-localize with collapsed actin structures. The mis-localization of keratin is interpreted as a result of the cells rounding up within the tissue because of a disruption in integrin-FN adhesive signaling.

Further supporting the disruption in PDGFr-α dependent signaling leading to a decrease in pFak at Tyr-397 and mesendoderm explants have larger focal adhesions. Larger focal adhesions are comparable to increased focal adhesion size found in FAK (-/-) mouse embryonic mesodermal cells (llić et al., 1995) or when Fak is knocked down in fibroblasts (Kim and Wirtz, 2013). Knocking down components required for integrin activation such as kindlin-2 or integrin β1 also resulted in an increase in focal adhesion size (Bandyopadhyay et al., 2012) similar to PDGFr-α knockdown. Taken together, these findings are consistent with integrin and PDGFr dependent signals acting cooperatively to modulate Fak phosphorylation at Tyr-397 and adhesion to FN necessary for focal adhesion assembly, directional protrusion formation, and cytoskeletal organization.

## Materials and Methods

### Collection of *Xenopus* eggs and embryos

*Xenopus* eggs were obtained fertilized in vitro. Embryos were allowed to develop and embryos were staged according to Nieuwkoop and Faber (Nieuwkoop and Faber, 1994). Embryos were dejellied in 2% cysteine, rinsed with dH20, and cultured in 0.1X MBS (MBS: 1X MBS: 88 mM NaCl, 1 mM KCl, 2.5 mM NaHCO_3_, 0.35 mM CaCl_2_, 0.5 mM MgSO_4_, 5 mM HEPES pH 7.8).

### Dorsal marginal zone explant preparation and single mesendoderm cell dissection

Glass coverslips were treated with 10N NaOH, rinsed successively in deionized water, 70% ethanol, 100% ethanol, and then flamed. Washed coverslips were coated with 10 μg/ml of bovine plasma FN (Calbiochem), or equimolar amounts of 9.11 (0.5 μM), 9.11a (0.5 μM) prepared in 1X MBS and left overnight at 4°C in a humidified chamber. FN coated coverslips were blocked with 0.45 μM sterile filtered 5% BSA. Mesendoderm was dissected and cells dissociated in Ca^2+^ and Mg^2+^ MBS. Mesendoderm cells were plated at sub-confluent density and single cells were blindly scored as round or spread (defined as 1 or more protrusion) (Ramos and DeSimone, 1996). Mesendoderm explants were prepared as described (Davidson et al., 2002; Weber et al., 2012). Briefly, the dorsal marginal zone of the *Xenopus* embryo was dissected at Stage 10^-^, secured with glass coverslips using silicon grease, and allowed to adhere for 1 hour to FN-coated glass coverslips.

### Morpholino knockdown, RNA transcription, and Biochemistry

Anti-sense morpholino oligonucleotides obtained from GeneTools (Philomath, OR) were used to knockdown *Xenopus laevis* PDGF-A Ligand (PDGF-A) and PDGF Receptor (PDGFr-α). All experiments were done with a total of 60ng morpholino injected per embryo. Sequences are as follows:

Control Morpholino: 5’- CCTCTTACCTCAGTTACAATTTATA-3’ (stock sequence)

PDGF-A Ligand Morpholino: 5′- AGAATCCAAGCCCAGATCCTCATTG-3′(Nagel et. al 2004)

PDGFrα Morpholino: 5’-GGCAGGCATCATGGACCGTAACAAC-3’

RNA encoding mCherry-LifeAct and GFP-XCK1(8) was transcribed from plasmid DNA and 5nLs were injected into the 2 dorsal blastomeres at the four cell stage for a final concentration of 500pg/ embryo. EGFP-XCK1(8) was obtained from V. Allan, University of Manchester.

### Western Blots

*Xenopus* embryos or dissociated mesendoderm tissue were solubilized in lysis buffer (100 mM NaCl, 50 mM Tris- HCl pH 7.5, 1% Triton X-100, 2 mM PMSF (phenylmethylsulphonylfluoride), protease inhibitor cocktail (Sigma)). Protein extracts were diluted with 2X Laemmli buffer (2% β- mercaptoethanol) and run on a 10% SDS-PAGE gel, transferred to nitrocellulose, and then probed using antibodies to pAkt T308 (1:1000, Cell Signaling #2965), pFak Tyr 397 (1:1000, Upstate), Total Akt (1:1000, Cell Signaling #3C67E7), and β-actin (1:10,000, Sigma #A3854).

### Immunofluorescence

Mesendoderm explants were fixed with ice-cold 100% methanol overnight at 4°C, rehydrated into TBS (75% methanol, 50% methanol, 25% methanol), washed with TBS-T, and stained overnight at 4°C with pan-cytokeratin C11 antibody (1:200, Sigma). After 3 washes with TBS-T, goat anti-mouse IgG conjugated to AlexaFluor488 was used to visualize cytokeratin. Actin was visualized using AlexaFluor488-actistain after mesendoderm explant fixation in 0.25% Glutaraldehyde, 3.7% Formaldehyde and 0.1% Tween20 for 10 mins at room temperature. Immunofluorescence imaging was performed on a Nikon C1 confocal microscope with a Nikon PlanApo/60×/1.40 objective.

### Tracking DMZ migration rates using ImageJ

Images were collected (1 per minute) using a Zeiss Axiovert 35 with OpenLab software (Improvision/Perkin Elmer, Waltham, MA). Image analysis was performed using Volocity and ImageJ (http://rsb.info.nih.gov; National Institutes of Health). Using ImageJ, each image was divided into left (0-341 pixels), middle (341-683 pixels), and right pixel (683-1024 pixels) sections. Tracing was done by placing 3 fiduciary marks (1 per section) along the leading edge of the mesendoderm explant using a 25 micron diameter paintbrush. In each case the x-coordinate was recorded and the average of the three dots was recorded as one data point. Explant retractions were defined as any movement that was opposite the direction of migration and recorded as negative values in distance traveled.

### Statistical Analysis

Graphs and statistical analysis were performed using PRISM V5 software. A Wilcoxon match-paired signed rank test was performed for a pairwise comparison when experiments had two treatment conditions with matched clutches of embryos. Differences between Control MO cells or PDGFr -α MO cells plated on 9.11 and 9.11a substrates were analyzed using a one-way ANOVA followed by Tukey’s post hoc test.

